# Spectral Signatures of L-DOPA-Induced Dyskinesia Depend on L-DOPA Dose and are Suppressed by Ketamine

**DOI:** 10.1101/2020.07.14.202721

**Authors:** Tony Ye, Mitchell J. Bartlett, Scott J. Sherman, Torsten Falk, Stephen L. Cowen

## Abstract

L-DOPA-induced dyskinesias (LID) are debilitating motor symptoms of dopamine-replacement therapy for Parkinson’s disease (PD) that emerge after years of L-DOPA treatment. While there is an abundance of research into the cellular and synaptic origins of LID, less is known about how LID impacts systems-level circuits and neural synchrony, how synchrony is affected by the dose and duration of L-DOPA exposure, or how potential novel treatments for LID, such as sub-anesthetic ketamine, alter this activity. Sub-anesthetic ketamine treatments have recently been shown to reduce LID, and ketamine is known to affect neural synchrony. To investigate these questions, we measured movement and local-field potential (LFP) activity from the motor cortex (M1) and the striatum of preclinical rodent models of PD and LID. In the first experiment, we investigated the effect of the LID priming procedures and L-DOPA dose on neural signatures of LID. Two common priming procedures were compared: a high-dose procedure that exposed unilateral 6-hydroxydopamine-lesioned rats to 12 mg/kg L-DOPA for 7 days, and a low-dose procedure that exposed rats to 7 mg/kg L-DOPA for 21 days. Consistent with reports from other groups, 12 mg/kg L-DOPA triggered LID and 80-Hz oscillations; however, these 80-Hz oscillations were not observed after 7 mg/kg administration despite clear evidence of LID, indicating that 80-Hz oscillations are not an exclusive signature of LID. We also found that weeks-long low-dose priming resulted in the emergence of non-oscillatory broadband gamma activity (> 30 Hz) in the striatum and theta-to-high-gamma cross-frequency coupling (CFC) in M1. In a second set of experiments, we investigated how ketamine exposure affects spectral signatures of low-dose L-DOPA priming. During each neural recording session, ketamine was delivered through 5 injections (20 mg/kg, *i*.*p*.) administered every 2 hours. We found that ketamine exposure suppressed striatal broadband gamma associated with LID but enhanced M1 broadband activity. We also found that M1 theta-to-high-gamma CFC associated with the LID on-state was suppressed by ketamine. These results suggest that ketamine’s therapeutic effects are region specific. Our findings also have clinical implications, as we are the first to report novel oscillatory signatures of the common low-dose LID priming procedure that more closely models dopamine replacement therapy in individuals with PD. We also identify neural correlates of the anti-dyskinetic activity of sub-anesthetic ketamine treatment.

## Introduction

Parkinson’s disease (PD) is a neurodegenerative disease with cardinal motor impairments of bradykinesia, rigidity, postural instability, and tremor (Olanow et al., 2009). These motor dysfunctions are caused by the death of dopaminergic neurons in the substantia nigra pars compacta (SNc) that project to the striatum, resulting in reduced dopaminergic tone in corticostriatal circuits. The gold-standard treatment for PD is dopamine replacement therapy via the dopamine precursor L-3,4-dihydroxyphenylalanine (L-DOPA). L-DOPA restores physiological dopamine concentrations in the striatum (Picconi et al., 2003) to reinstate voluntary motor activity. However, prolonged L-DOPA exposure eventually leads to incapacitating L-DOPA-induced dyskinesias (LID) (Cotzias et al., 1969), making untenable continued L-DOPA treatment. Consequently, new approaches are needed for treating LID and extending L-DOPA’s window of clinical efficacy.

Neuroplastic changes associated with prolonged L-DOPA are believed to alter circuits in the basal ganglia and motor cortex (M1), and these alterations result in beta (∼20 Hz) and high-gamma (∼80 Hz) oscillations (Litvak et al., 2011) that are evident in patients with LID. High gamma may be a unique signature LID as clinical studies with human Parkinson’s disease patients experiencing LID have identified 80-Hz activity in the subthalamic nucleus (STN) (Alonso-Frech et al., 2006; Lopez-Azcarate et al., 2010; Swann et al., 2016; Williams et al., 2002), 70-80 Hz spectral power and coherence from surface electroencephalographic (EEG) probes (Williams et al., 2002), and ∼80-Hz oscillations and coherence from cortical surface electrodes placed over motor cortex and depth electrodes in the STN (Swann et al., 2016).

Similar 80-Hz gamma activity has been identified in preclinical rodent models of LID (Dupre et al., 2016; Halje et al., 2012). For example, several groups have utilized a high-dose 7-d L-DOPA priming protocol (12 mg/kg) in 6-hydroxydopamine-(6-OHDA) lesioned animals and found that 12 mg/kg L-DOPA induces clear narrow-band 80-Hz gamma in M1. This 80-Hz oscillation is sometimes referred to as “finely-tuned gamma”, and is associated with abnormal involuntary movements (AIMs) produced by L-DOPA in animal models (Dupre et al., 2016; Halje et al., 2012). The presence of a clear 80-Hz signature of LID in human subjects and animal models suggest that treatments that suppress mechanisms that produce corticostriatal 80-Hz gamma may reduce LID symptoms.

Focal 80-Hz gamma in LID differs from less focal low, medium, and high gamma activity associated with facilitating neuronal communication (Buzsáki, 2010; Lisman and Idiart, 1995; Lisman and Jensen, 2013), plasticity (Buzsáki and Wang, 2012; Colgin et al., 2009; Galuske et al., 2019), and motor activity (Johnson et al., 2017; Muthukumaraswamy, 2010). Focal 80-Hz gamma also differs from desynchronized broadband gamma (∼30-80 Hz) that is associated with increased excitability in neural circuits (Sohal and Rubenstein, 2019; Yizhar et al., 2011) and uncorrelated synaptic and spiking activity (Belluscio et al., 2012). Indeed, broadband gamma can be evoked through disinhibition of principal cells by reducing glutamatergic drive to parvalbumin-expressing inhibitory interneurons through genetic manipulations (Cho et al., 2015; delPino et al., 2013) and delivery of NMDAR antagonists such as ketamine or MK-801 (Caixeta et al., 2013; Carlén et al., 2012; Hakami et al., 2009; Hudson et al., 2020; Kulikova et al., 2012; McNally et al., 2020; Pinault, 2008; Rivolta et al., 2015; Ye et al., 2018).

Most physiological studies of LID have focused on the motor cortex, STN, and dorsal striatum. As a result, little is known about how LID affects neural activity in the NAc. This is surprising as shrinkage and dopamine depletion of the NAc occurs at later stages of PD, and these changes are associated with depression and cognitive deficits in PD patients (Hanganu et al., 2014; Mavridis et al., 2011). Cognitive impairment in PD likely involves mesocorticolimbic dopamine dysfunction. For example, PD patients on-but not off-medication are impaired at probabilistic reversal learning (Cools et al., 2001; Torta et al., 2009), a task involving the NAc (Dalton et al., 2014). Furthermore, L-DOPA therapy suppresses activity in the NAc but not dorsolateral striatum or prefrontal cortex, which, in turn, biases probabilistic reversal learning (Cools et al., 2007). Depression affects >30% of PD patients (Slaughter et al., 2001), and this likely results from reduced dopaminergic transmission in the NAc (Nestler and Carlezon, 2006). Interestingly, ketamine exerts a dose- and time-dependent effect on NAc oscillatory activity that may play a role in the drug’s antidepressant effect (Manduca et al., 2020). Thus, even though dopamine depletion in PD is less severe in the NAc than in the dorsal striatum, reduced NAc dopamine is likely involved in the cognitive and affective symptoms of PD.

A complication of work with animal models of dyskinesia is the common use of multiple priming protocols to induce LID (Bartlett et al., 2016; Cenci et al., 1998; Dekundy et al., 2007; Dupre et al., 2016; Halje et al., 2012). Consequently, it is possible that some of the identified oscillatory signatures of LID in animal models are unique to the dosage and duration of L-DOPA exposure. Broadly, LID priming procedures can be divided into low-dose+long-duration (*e*.*g*., 6-7 mg/kg for 21 d; Cenci et al., 1998; Dekundy et al., 2007; Bartlett et al., 2016) and high-dose+short-duration protocols (*e*.*g*., 12 mg/kg, 7-d; Halje et al., 2012*b*; Dupre et al., 2016). There are clear advantages to both procedures. For example, the high-dose priming results in the LID behavioral phenotype on the first day of exposure, which can be of significant practical advantage (Carta et al., 2006). In contrast, low-dose (7 mg/kg) 21-d priming produces more gradual development of LID that better reflects the gradual development of LID in patients. No study to date has investigated the influence of priming protocol and L-DOPA dosage on oscillatory signatures of LID. Determining whether LID-associated oscillations are impacted by the dosage or duration of L-DOPA administration is important given the variety of clinical dosages of L-DOPA given to PD patients. In addition, understanding how L-DOPA dosage and exposure duration impacts LID-associated oscillatory activity could reveal mechanisms underlying LID generation and development.

Low-dose sub-anesthetic ketamine has been successfully used to treat chronic pain (Niesters et al., 2014) and treatment-resistant depression (Andrade, 2017; Diamond et al., 2014), with *S*-ketamine now being an FDA approved drug for use in treatment-resistant depression (Kaufman, 2019). A retrospective case study in PD patients receiving sub-anesthetic infusions of ketamine to treat pain showed reduced LID for up to one month (Sherman et al., 2016). Ketamine treatment can also reduce LID long-term in rodent models of established LID and of LID development (Bartlett et al., 2020, 2016). These lasting effects may be due to ketamine’s ability to modify oscillatory activity throughout the brain (Hunt and Kasicki, 2013; Ye et al., 2018). A single injection of sub-anesthetic ketamine is known to trigger oscillatory activity throughout corticostriatal and hippocampal circuits (Caixeta et al., 2013; Nicolás et al., 2011; Olszewski et al., 2013). In addition, we have shown in naïve rats that repeated sub-anesthetic injections trigger region-wide and dose-dependent high-frequency oscillations (HFO, >100 Hz), broadband asynchronous gamma (40–80 Hz), and cross-frequency interactions (Ye et al., 2018).

In this study, we investigated whether the LID-induction protocol or the L-DOPA dose administered during LID induction resulted in unique oscillatory signatures. We also investigated the hypothesis that ketamine produces its anti-dyskinetic effect by suppressing neural oscillations associated with LID. These questions were investigated through the measurement of local-field activity in M1 and dorsolateral and dorsomedial striatum (DLS, DMS), regions most often associated with motor deficits in PD and LID. Measurements were also acquired from the nucleus accumbens (NAc) given its strong interconnectivity with the dorsal striatum and M1, as well as the association between NAc dysfunction and cognitive and affective deficits in PD (Hanganu et al., 2014; Mavridis et al., 2011). All measurements were acquired from awake and behaving unilateral 6-OHDA-lesioned rats treated with L-DOPA.

## Materials and Methods

### Animals

Twenty-eight male Sprague-Dawley rats (6 weeks old, 250-275 g at arrival, Harlan Laboratories, Indianapolis, IN) were single-housed in a temperature and humidity controlled 12-hr reverse light/dark cycle room with food and water available *ad libitum*. Rats were divided into four groups based on lesion or L-DOPA treatment: naïve non-lesioned controls (*n*=8), PD (*n*=7), LID 7-d priming (*n*=6), and LID 21-d priming (*n*=7). Sample sizes for groups were determined by previous experiments (Ye et al., 2018). Following the initial set of experiments (days 1-10), animals in the 7-d priming group were kept in the colony room and did not receive any L-DOPA priming injections for 28 days (**Fig. 1A**). This condition allowed us to investigate what occurs when the effects of priming wears off. All procedures were in accordance with NIH guidelines for the Care and Use of Laboratory Animals and approved IACUC protocols at the University of Arizona. The experimental design for the studies is summarized in **Fig. 1**.

**Figure 1.**
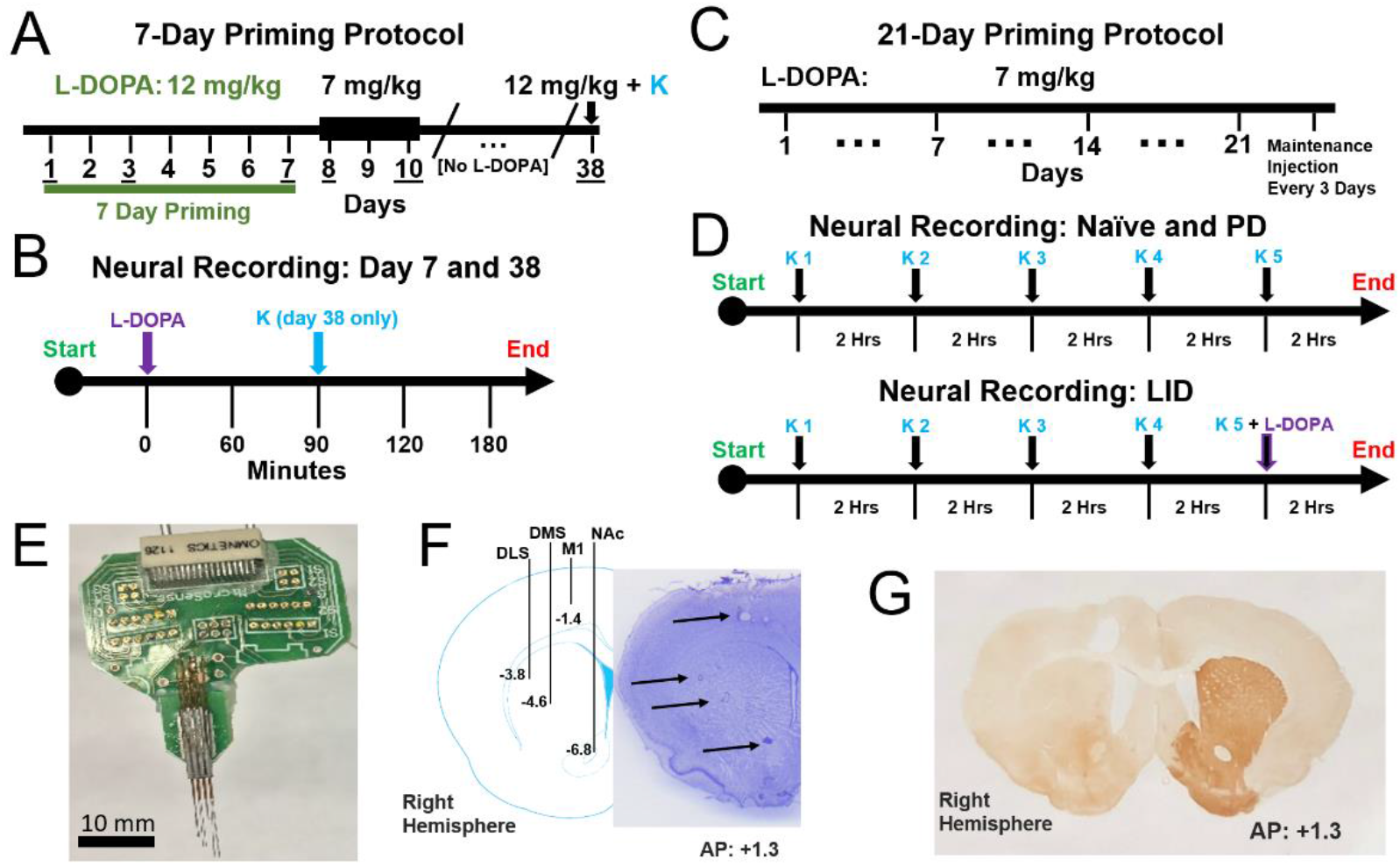
Experimental Design and Neural Recordings. **(A)** Schematic of 7-day priming protocol in Experiment 6-OHDA-lesioned animals received L-DOPA (*i*.*p*., 12 mg/kg) for 7 consecutive days. Animals then received a 7 mg/kg injection of L-DOPA on days 8-10. Following a 28-day abstinence from L-DOPA, the same animals once again received a single injection of L-DOPA (12 mg/kg) paired with ketamine (20 mg/kg). **(B)** Timeline of neural recording sessions on days 7 and 38 of the 7-day LID priming animals. Recordings began with 1-hr baseline. Animals were then injected with L-DOPA (12 mg/kg). LAO + AIMs scored every 20 minutes. On Day 38 (but not Day 7) ketamine (20 mg/kg, *i*.*p*.) given 90 minutes after L-DOPA. **(C)** Schematic of 21-day priming protocol for Experiment 6-OHDA-lesioned animals were administered L-DOPA (7 mg/kg) daily for 21 days. After 21 days, maintenance injections of L-DOPA (7 mg/kg) were given every 3 days. Neural recordings were not conducted during the 21-day priming period. **(D)** Timeline of neural recordings for naïve and PD animals (top) and 21-day priming LID animals (bottom). Recordings began with a 1 hr baseline followed by a ketamine injection (20 mg/kg, *i*.*p*.) every two hours, totaling to 5 injections. For LID animals (bottom), the 5^th^ ketamine injection was paired with L-DOPA (7 mg/kg). A custom-made 32-channel electrode surgically implanted into the right hemisphere of all experimental animals. Schematic of electrode array placement (AP: +1.3, ML: +2.7 centered, DV: −6.8 deepest electrode) and representative example of histological verification of targets. RIGHT: Verification of 6-OHDA lesioning in 6-OHDA and LID animals via tyrosine-hydroxylase staining. Expression of tyrosine-hydroxylase (dark pigmentation; left hemisphere) is a marker of functioning dopaminergic neurons. **(G)** Verification of a successful (>90%) 6-OHDA lesion results in a light pigmentation of the striatum (right hemisphere). Behavioral verification of 6-OHDA lesion

### Unilateral 6-OHDA-lesion PD model

As previously published (Bartlett et al., 2016), 6-OHDA hydrochloride was injected into 2 locations (5 μg/site) of the medial forebrain bundle: MFB: AP = −1.8 mm, ML= +2.0 mm, DV= −8.2 mm and AP = −2.8 mm, ML= +1.8 mm, DV = −8.2 mm. Amphetamine-induced (5.0 mg/kg, *i*.*p*., Sigma-Aldrich) rotations were administered ∼3 weeks post-surgery and were scored by blinded experimenters to assess the degree of lesion (**Supplementary Table 1**). An average score ≥5 corresponds to >90% dopamine depletion (Dekundy et al., 2007). Rats that reached criteria were divided into three groups: LID induction with 7-d priming (*n*=6), 21-d priming (*n*=7), or PD (*n*=7).

### Electrode implantation

Following the procedure reported in Ye et al., 2018, rats were anesthetized with isoflurane and implanted with two custom-made 32-channel electrode arrays by the experimenters (**Fig. 1E**), with each array composed of 16 twisted-wire stereotrodes (California Fine Wire Co., Grover Beach, CA). All recordings were referenced to a cerebellar skull screw. The anterior array was placed in the right hemisphere, and individual stereotrodes targeted M1 (AP:+1.3, ML:+2.3, DV:-1.4), DLS (AP:+1.3, ML:+3.5, DV:-3.8), DMS (AP:+1.3, ML:+2.9, DV:-4.6), and NAc (AP:+1.3, ML:+1.7, DV:-6.8) (**Fig. 1F**). The posterior array was implanted over the hippocampus (centered at AP: −3.0, ML: +2.2) with electrodes lowered near the fissure (DV: −3.2), CA1 (DV: −2.3), dentate gyrus (DV: −3.8), and S1 (DV: −1.4). Data from the posterior hippocampal array was not included in analyses. Rodent welfare was monitored daily by experimenters and animal care staff. Post-operative analgesia: 5 mg/kg (*s*.*c*.) Carprofen (Zoetis, Parsippany, NJ) for 48 h post-surgery. Topical anti-biotic ointment (Water-Jel Technologies, Carlstadt, NJ) given for up to 5 days as needed. The non-lesioned naïve control rats were surgically implanted 2-3 weeks after arrival (∼2 months old). The PD and LID 7 d priming groups were implanted approximately one week after amphetamine rotation tests (∼5 months old), and the LID 21-d priming group were implanted after the 21-d priming period (∼6 months old).

### Drug treatments

Using a paradigm described in Bartlett and colleagues (2016), drugs were delivered during neural recording sessions through five intraperitoneal (*i*.*p*.) injections of ketamine or saline (**Fig. 1D**). The first injection was delivered (6 AM) after one hour of baseline recording. Each injection was separated by two hours. Injections were either ketamine hydrochloride (20 mg/kg) (Clipper Distributing, St. Joseph, MO) or 0.9% saline (SAL) solution. For the rats in the LID-priming group (LID rats), the 5^th^ injection of ketamine or SAL included a co-injection of L-DOPA (7 mg/kg, *i*.*p*., Sigma-Aldrich).

### 7-d and 21-d LID priming in 6-OHDA-lesioned rats, maintenance, and behavioral analysis

Approximately 6 weeks after the 6-OHDA lesion, rats in the high-dose 7-d L-DOPA priming group (*n*=6) were treated daily with L-DOPA (12 mg/kg) + benserazide (15 mg/kg, *s*.*c*., Sigma-Aldrich) for 7 d (*i*.*e*., during L-DOPA priming) (**Fig. 2A-D**). Rats in the low-dose 21-d L-DOPA priming group were treated daily with L-DOPA (7 mg/kg, *i*.*p*.) + benserazide (14 mg/kg) for 3 weeks, as previously published (Bartlett et al., 2016) (**Fig. 1C**). Rats in the 21-d group received maintenance doses of L-DOPA (7 mg/kg + 15 mg/kg benserazide) following the 21-d priming period (two doses per week, every 2-3 days for the remainder of the experiment). L-DOPA-induced AIMs were scored by an experimentally blinded investigator on a scale from 0 to 4, and cumulative limb, axial, and orolingual (LAO) scores for all LID rats across all sessions are presented in **Fig. 3B** as previously published (Bartlett et al., 2016). After the 21-d priming period, rats that met behavioral criteria (*n*=7) proceeded to surgical implantation, and were given maintenance injections of L-DOPA (7 mg/kg) every 2-3 days to maintain stable dyskinesia (Bartlett et al., 2016). This contrasts with the 7-d priming group that was implanted prior to L-DOPA exposure. Neural recordings began one week following implantation.

**Figure 2.**
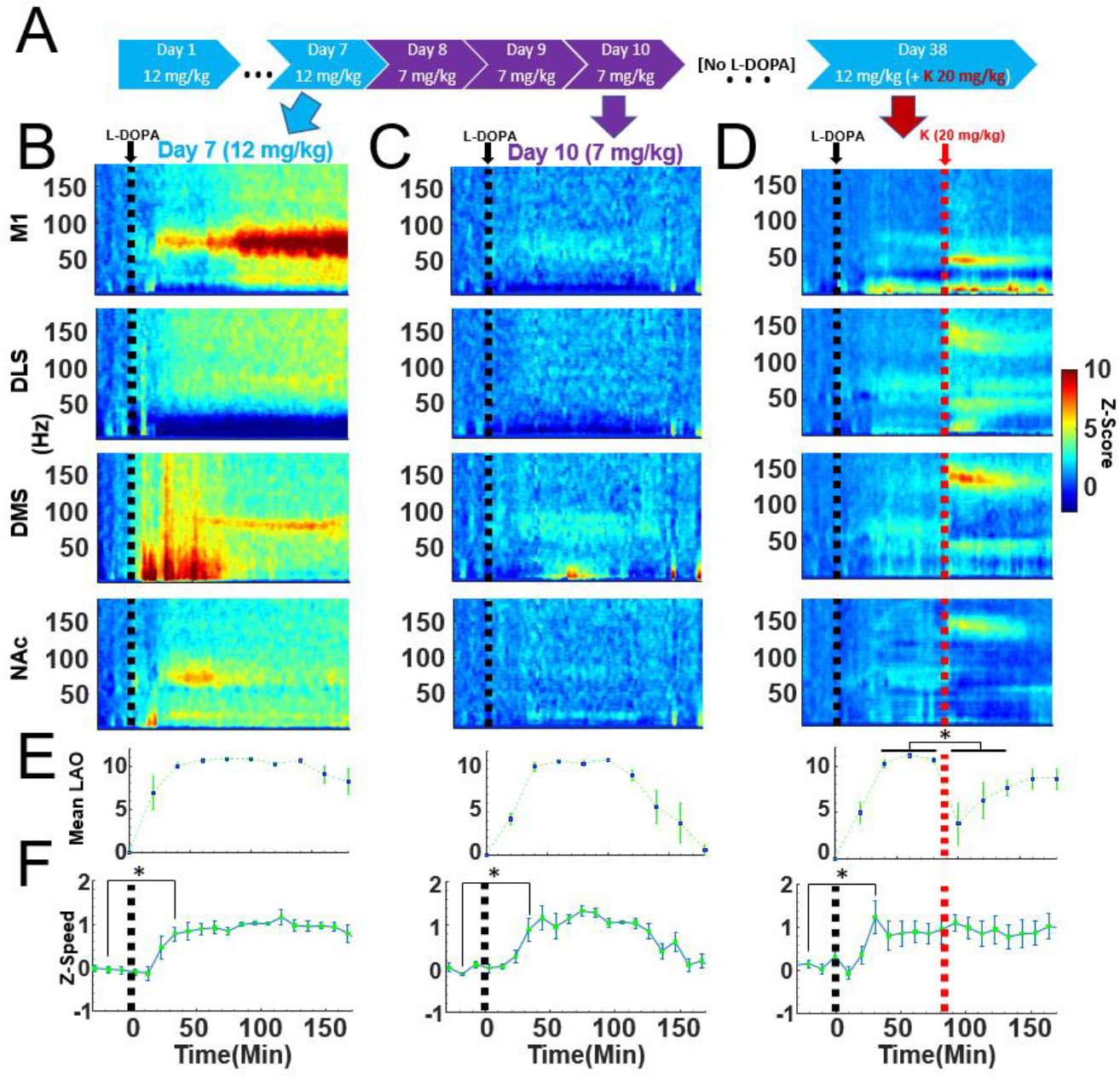
Experiment 1: Priming dose- and duration-dependent oscillatory responses of L-DOPA in LID animals. Timeline of 7 d L-DOPA priming (12 mg/kg) and subsequent injections (as in Fig. 1A). Arrows indicate the neural recording days used for B-D. **(B)** Average spectral power (*n*=6) of the 7 d L-DOPA priming protocol on the 7^th^ day of exposure in M1, DLS, DMS, and NAc *(n*=6). All neural activity normalized to pre-injection baseline (−32 to −2 min). **(C)** After the 7^th^ day of 12 mg/kg, the same animals received 7 mg/kg on Days 8-10. Average spectral power is shown for Day 10 (7 mg/kg). **(D)** After abstaining from L-DOPA for 28 days, animals once again received 12 mg/kg of L-DOPA followed 90 minutes by an injection of ketamine (20 mg/kg). **(E)** Cumulative Limb, Axial, and Orolingual (LAO) scores (mean±SEM) for each 20 min time point of the recording sessions on Days 7, 10, and 38. Day 38 (right): Ketamine significantly reduced LAO scores (40 minutes before vs 40 minutes after ketamine injection, *t*-test, *p*=0.02, *d*=2.0). **(F)** Gross locomotor activity (mean±SEM) significantly increased 10 min before L-DOPA vs 40 min after for Day 7, 10, and 38 (*t*-test, all *p<*0.05).

**Figure 3.**
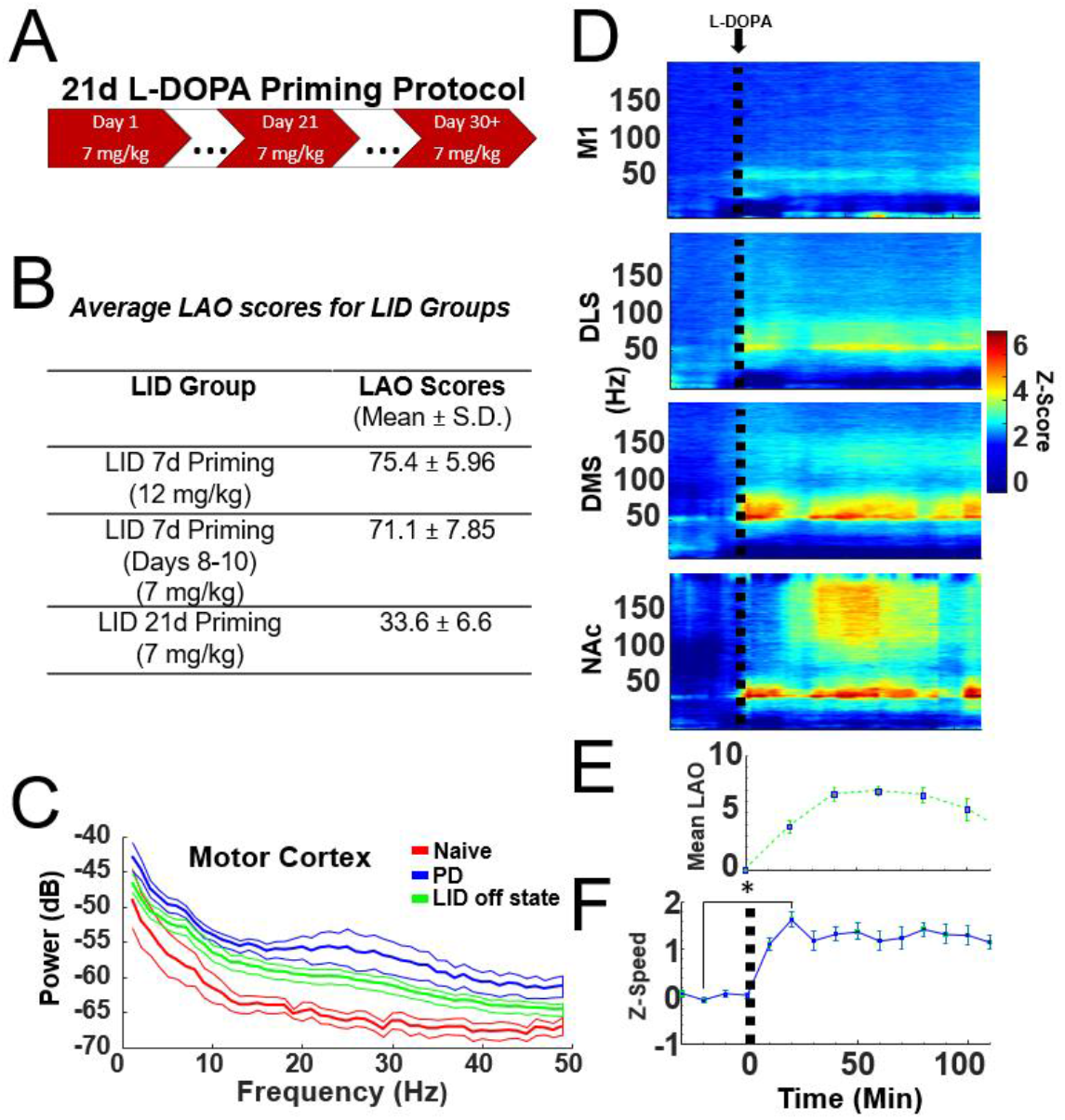
LID animals primed for 21 days displayed a different spectral signature than 7-day priming. **(A)** Timeline of 21-day L-DOPA priming protocol. 6-OHDA-lesioned animals received daily injections of L-DOPA (7 mg/kg) for 21 days. After this priming period, animals continued to receive 7 mg/kg of L-DOPA every 3 days to maintain stable dyskinesia. **(B)** Average (Mean ± S.D.) cumulative LAO AIMs across all sessions in LID animals in 21- and 7-day priming groups (during the 180 mins testing). After the 7-day (12 mg/kg) priming, the same group of animals then received L-DOPA at 7 mg/kg on days 8-10. **(C)** Average baseline (−32 to −2 min prior to L-DOPA injection) power (Mean ±SEM) of Naïve (*n*=8), PD (*n*=7), and LID (off drug; *n*=7) animals. Mean ± SEM. **(D)** Average spectral power surrounding a L-DOPA injection (7 mg/kg, post 21-day priming). **(E)** Cumulative Limb, Axial, and Orolingual (LAO) scores (mean±SEM) for each 20 min time point on the final day of the 21-d LID priming, prior to the neural recording sessions of (D). **(F)** Average locomotor activity (mean±SEM) significantly increased 10 min before L-DOPA vs. 40 min after (*t*-test, *p=*0.02, n=7).

### Neurophysiological recordings

A 256-channel data acquisition system (KJE-1001, Amplipex Ltd.) was used for neural recordings. A light-emitting diode (LED) was attached to the rat’s implant for video tracking (Manta G-033C, Allied Vision, Exton, PA). Recordings and drug injections were conducted in a polycarbonate cage (47cm x 51cm x 20cm) once per week for each animal commencing at 5 AM. Food and water were available *ad libitum*. Recordings for the 7-d priming experiments were conducted in a large (150cm x 150cm x 23cm) open field to limit artifacts from dyskinetic animals hitting cage walls. AIMs were scored during each neurophysiological recording only for the animals in the 7-d priming group (**Fig. 1A**).

### Histology and immunohistochemistry

Direct current stimulation (20 µA for 20 s) was used for electrolytic lesions at each recording site. Three-days following the lesion, rats were injected with a fatal dose of Euthasol (0.35 mg/kg, *i*.*p*.; Virbac, Fort Worth, TX) and transcardially perfused via phosphate buffered saline and 4% paraformaldehyde. A frozen microtome was used to produce coronal sections (40 µM) for Nissl staining (Ye et al., 2018) (**Fig. 1F**) and tyrosine hydroxylase (TH; Bartlett et al., 2016) (**Fig. 1G**) for verification of electrode placement and dopamine-depleted striatum, respectively.

### Data pre-processing and statistical analysis

Raw LFP signals were acquired at 20 kHz and down sampled to 500 Hz for analysis. Absolute values of the LFP trace that exceeded 1.5 mV or the 99.98^th^ percentile after cross-band power (2-160 Hz) summation were considered artifact and omitted from the analyses. To reduce the impact of volume-conduction, signals were locally re-referenced using a second within-region electrode (0.7 mm inter-electrode distance) at the same depth (Ye et al., 2018). ANOVAs and Student’s *t*-tests (α=0.05) were used to assess statistical significance. All *post-hoc* comparisons were Tukey-Kramer or Holm corrected to adjust *p*-values. All analyses were performed using MATLAB.

### Analysis of spectral activity, cross-frequency coupling, and movement speed

The spectral power across frequency bands was determined using a fast Fourier transform spectrogram (frequency bin=0.5 Hz, 10 s Hanning window, spectrogram() in Matlab). The frequency bands used for statistical analysis were defined as follows: delta (1–4 Hz), theta (5–10 Hz), beta (15–30 Hz), low-gamma (35–55 Hz), high-gamma (70–85 Hz), broadband gamma (35-–85 Hz), and HFO (120–160 Hz). Gamma was divided into separate bands given evidence for distinct roles played by different frequencies in neuronal communication (Buzsáki, 2010; Lisman and Idiart, 1995; Lisman and Jensen, 2013), plasticity (Buzsáki and Wang, 2012; Colgin et al., 2009; Galuske et al., 2019), and motor activity (Johnson et al., 2017; Muthukumaraswamy, 2010). Broadband gamma was also included given the association between broadband activity and increased neuronal excitability and desynchronized activity (Sohal and Rubenstein, 2019; Yizhar et al., 2011). The band used here to define high gamma is narrower than some definitions given the focal nature of 80-Hz activity in LID.

To address the issue of power-law 1/f scaling, data was normalized using the Z-transform (Cohen, 2014). The baseline mean (−32 to −2 min preceding the first injection) was subtracted from the spectral power and then divided by the standard deviation (S.D.) to yield a z-score. Video tracking of rat’s movement was measured in cm/second (s). This data was also normalized using the baseline mean with same Z-transform procedure to yield a z-score (i.e., Z-speed) to assess change in overall movement compared to the baseline period.

Phase-amplitude cross-frequency coupling (PAC) was measured as described in Cohen (2014) and Ye et al., (2018). First, LFP signals were filtered in the target low- and high-frequency bands using a Butterworth filter (fs=500 Hz, order=6). Phase was extracted using a Hilbert transform. Power was extracted as the envelope of the absolute value of the filtered signal. CFC was computed as 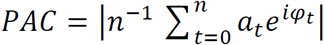 where *a* is high-frequency power and ϕ is the phase of the low-frequency signal. This value was compared to values computed using a randomized shuffle control (*n*=200 permutations). The mean and SD of this null hypothesis distribution were used to convert the measured PAC score into a z score (PACz). All data in figures show mean ± SEM unless otherwise noted.

### Data Availability

The data are available from the corresponding author upon reasonable request.

## Results

### Dyskinesias and oscillatory activity in PD and LID model animals

Beta oscillations (15–30 Hz) in M1 are a signature of PD. We examined baseline oscillatory activity in M1 of PD, LID (off-state), and naïve control animals (**Fig. 3C**). As expected, significant differences in beta power were observed (ANOVA, *F*(2,18)=7.71, *p*=0.004, η^2^=0.25). *Post-hoc* comparisons revealed that the PD (*p*=0.002, *n*=7) and LID groups (*p*=0.04, *n*=7) had greater beta power than naïve controls (*n*=8) (**Supplementary Fig. 1**). Beta power did not differ between PD and LID animals (*p*=0.11). Amphetamine-induced rotations were observed in these animals (>5 rotations/min; **Supplementary Table 1**), supporting the behavioral PD phenotype. Immunohistochemical analysis verified dopamine-depletion in the PD and LID groups (**Fig. 1G**).

The 6-OHDA-lesioned animals that met rotation criteria were assigned to either PD, LID 7 d priming, or LID 21 d priming groups. Cumulative limb, axial, and orolingual (LAO) scores during the L-DOPA on-state clearly indicated the LID phenotype in the 7 d (12 mg/kg) (75.4±5.96; Mean±S.D.), post 7 d (7 mg/kg) (71.1±7.85), and 21 d (7 mg/kg) (33.6±6.6) groups (**Fig. 3B**). LAO scores were consistent with previous literature (Bartlett et al., 2016; Dupre et al., 2016; Halje et al., 2012).

### Narrow-band 80-Hz high-gamma depends on L-DOPA dose

LID is associated with a narrow-band 80-Hz activity in M1 and DLS which is correlated with AIMs onset and duration (Halje *et al*., 2012; Dupre *et al*., 2016). Similarly, we observed narrow-band 80-Hz oscillations in M1, DMS, and NAc on the 7^th^ day of high-dose L-DOPA priming (**Fig. 2B**) following 12 mg/kg L-DOPA LID induction, but not on day 10 when 7 mg/kg L-DOPA was administered (**Fig. 2C**). Both 12 mg/kg and 7 mg/kg doses resulted in expression of clear LID (**Fig. 2E**) with significant increases in overall movement before vs. after L-DOPA injection (*t*-test, all *p*<0.05) (**Fig. 2F**). Statistical comparisons were performed by taking mean high-gamma power (dB) in the 22-60 min post-L-DOPA window given that L-DOPA requires ∼20 minutes to become active in the CNS. ANOVA and post-hoc analyses identified a strong increase in narrow-band high-gamma in M1 (**Fig. 4A**; ANOVA, *F*(4,28)=12.38, *p*=0.0001, η^2^=0.42), and smaller main effects in DMS (*F*(4,28)=6.56, *p*=0.03, η^2^=0.16) and NAc (*F*(4,28)=8.53, *p*=0.002, η^2^=0.39) on Day 7 of 12 mg/kg priming. The increase in narrow-band 80-Hz power was only identified when 12 mg/kg L-DOPA was used to induce LID. This is the first report to our knowledge of LID-associated 80-Hz power in the DMS. Furthermore, these data indicate that lower doses of L-DOPA induce strong LID behaviors in the absence of narrow-band 80-Hz oscillations.

**Figure 4.**
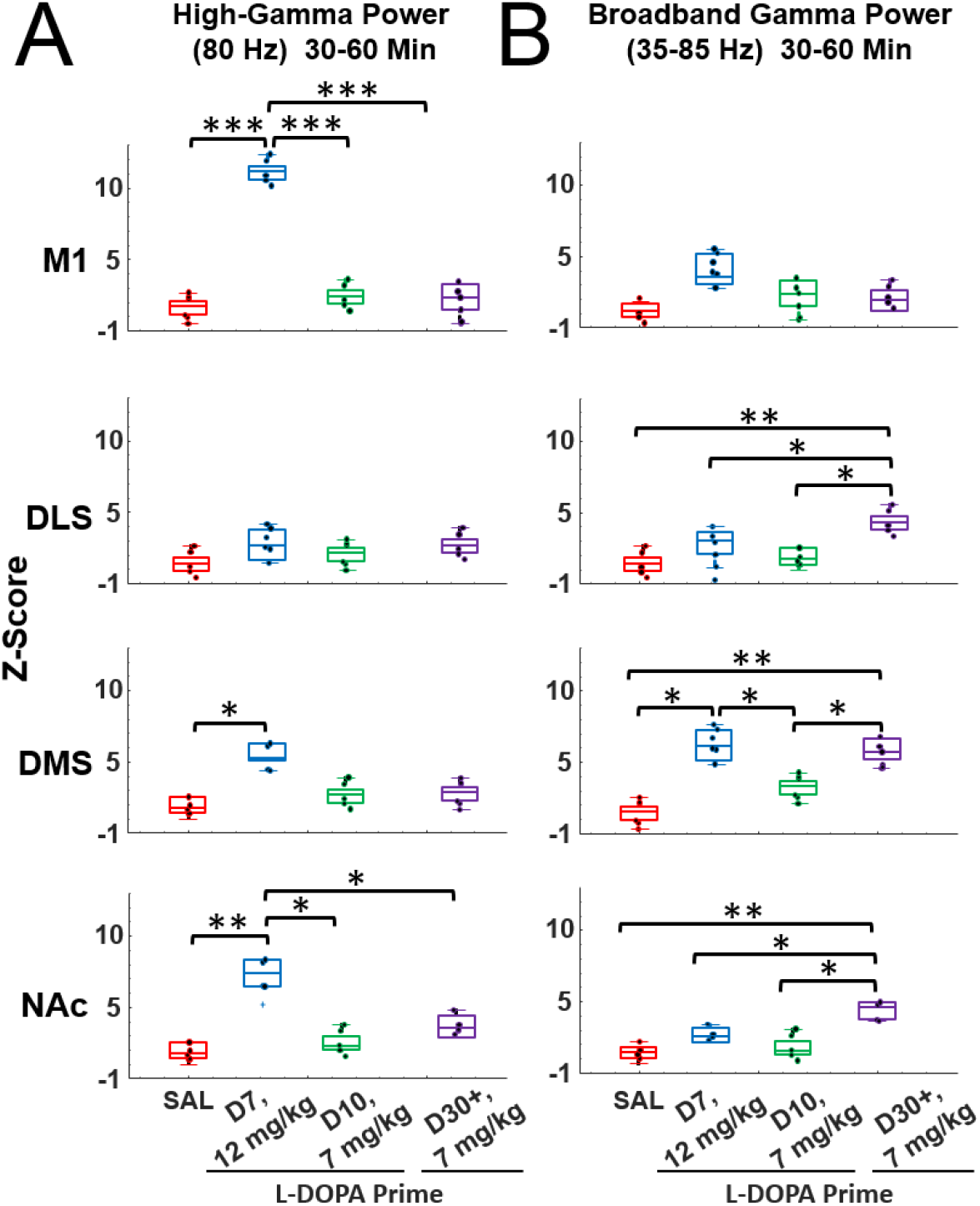
Priming- and dose-dependent gamma activity after L-DOPA administration in 6-OHDA-lesioned animals. **(A)** Data from Days 7, 10, 38 (Fig. 2), and post-21-d priming (Fig. 3D). High-gamma (80 Hz) activity in M1, DLS, DMS, and NAc during the 30-60 min post-L-DOPA injection period for saline (red), 7 d 12 mg/kg (blue), post-7 d 7 mg/kg (green), and 21 d 7 mg/kg (purple) conditions. L-DOPA (12 mg/kg, *n*=6) triggered significant increases in 80 Hz high-gamma compared to all conditions in the M1 and NAc (ANOVA; all *p*<0.05, Tukey-corrected). High-gamma was only significantly greater than SAL in the DMS (ANOVA; *p*=0.03, Tukey-corrected). As in (A) but for broadband gamma (35-85 Hz). Broadband gamma was significantly greater after the 21 d L-DOPA (7 mg/kg, *n*=7) priming in the DLS and NAc compared to 7-d priming and SAL (ANOVA; all *p*<0.05, Tukey-corrected). However, broadband gamma in the DMS in Day 7 of 12 mg/kg priming was not statistically different compared to 21-d priming animals. These data suggest differential oscillatory signatures of L-DOPA in LID animals that are dose- and priming-duration-dependent.

### Broadband gamma emerges after weeks-long exposure to low-dose L-DOPA

While injections of high-dose (12 mg/kg) L-DOPA induced robust narrow-band 80-Hz gamma on Day 7 of priming, low-dose (7 mg/kg) L-DOPA delivered on Day 10 did not (**Fig. 2C**).

However, after 30+ days of L-DOPA exposure in a separate group of animals, the same low-dose 7 mg/kg injection of L-DOPA appeared to increase broadband gamma power (>30 Hz) in the striatum, but not M1 (**Fig. 3D**). Although we did not score LAO during these neural recordings, the cumulative LAO taken on the final day of the 21-d priming procedure indicate dyskinesia indeed developed (**Fig. 3E**), and despite the lack of narrow-band gamma, a significant increase in overall locomotor activity was also observed during these neural recordings (*t*-test, *p*=0.02. **Fig. 3F**). To investigate the potential differences in gamma, we defined broadband gamma as gamma power between 35-85 Hz and assessed mean broadband activity after at least 21 days of priming (see **Supplementary Fig. 2** for a comparative analysis of low and high gamma power). Analysis of broadband gamma in the 21+ day condition during the L-DOPA on-state revealed that broad band power increased in the NAc, DMS, and DLS but not in M1 (**Fig. 4B**, *F*_*DLS*_(4,28)=8.53, *p*=0.002, η^2^=0.23; *F*_*DMS*_(4,28)=10.22, *p*=0.001, η^2^=0.35; *F*_*NAc*_(4,28)=6.31, *p*=0.02, η^2^=0.15; *F*_*M1*_(4,28)=2.46, *p*=0.78, η^2^=0.01). *Post-hoc* analyses within each striatal region indicated that this effect was present after at least 21 days of priming (Tukey corrected *post-hoc* comparisons between SAL and 21-d, 7 mg/kg were significant at *p*<0.01 for DLS, DMS, and NAc), suggesting that long-duration L-DOPA exposure enhanced broadband gamma activity. We also observed significant increases in broadband gamma in the DMS during Day 7 of L-DOPA (12 mg/kg) in the 7-d priming group compared to SAL (*p*=0.01) and Day 10 (*p*=0.03) of 7 mg/kg. Despite this observation, broadband gamma did not coincide simultaneously with narrow band gamma in the DMS. These data indicate that unique spectral signatures of LID emerge as a function of the dose used during LID induction and the duration of the priming.

### Exposure to ketamine in LID animals reduces striatal gamma activity but increases M1 gamma during the L-DOPA on-state

The acute effect of ketamine injection on ongoing spectral activity in LID animals was investigated since ketamine exposure reduces AIMs scores in LID animals (Bartlett et al., 2016). LID was induced in rats using the low-dose+long-duration LID induction procedure (7 mg/kg, 21 d to establish stable and moderate LID and a cumulative LAO score of 33.6±6.6, Mean±S.D., **Fig. 3B**). These animals received a 10-hour exposure to ketamine (**Fig. 1D**) as this was the exposure protocol determined to reduce LID (Bartlett et al., 2016). Spectral activity surrounding each of the 5 ketamine injections was visualized using baseline-normalized spectrograms (**Fig. 5A**). Baseline was defined as the mean spectral power measured during the −30 to −2 min interval preceding Injection 1. The mean and S.D. of this activity was used to generate a z-score measure of power. The time-course of activity in each of the targeted frequency bands surrounding Injection 1 (ketamine alone) and 5 (L-DOPA+ketamine) is presented in **Fig. 5B**.

**Figure 5.**
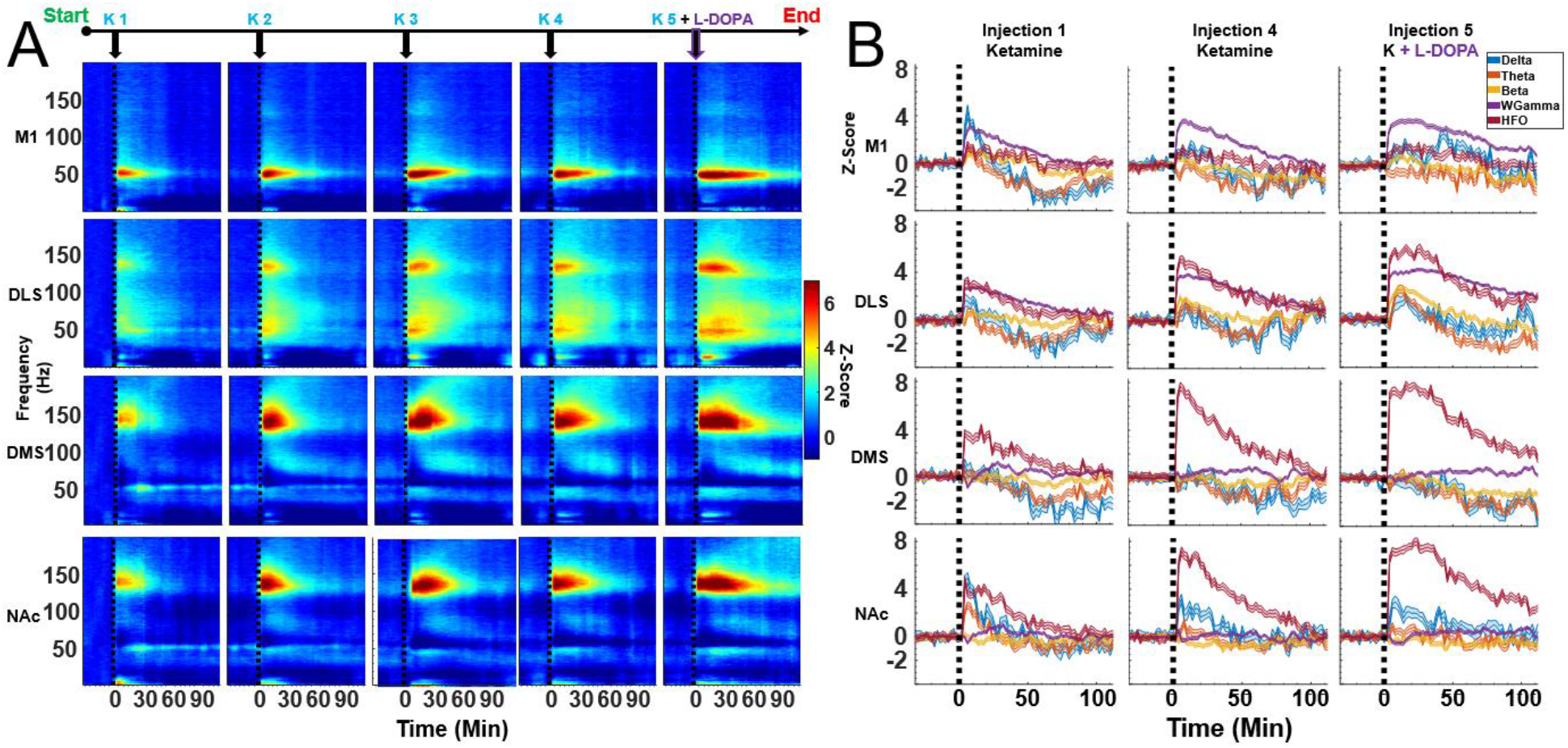
Average spectral responses to ketamine and ketamine + L-DOPA injection in 21-d LID animals. **(A)** Responses were normalized to the pre-injection baseline (−32 to −2 mins before 1^st^ injection). All data are from 21-d LID primed animals *(n*=7) recorded between 30-60 days post-priming. Single injections of ketamine (20 mg/kg) were administered every two hours. The 5^th^ and final injection were co-administration of ketamine + L-DOPA (7 mg/kg) to trigger the L-DOPA on-state. **(B)** Time course of spectral responses following ketamine injections 1, 4, and 5 (ketamine + L-DOPA) by frequency band (Mean±SEM).

We tested the hypothesis that L-DOPA+ketamine reduces striatal broadband gamma activity that was associated with LID under the low-dose+long-duration procedure (**Figs. 3, 4**). This hypothesis was explored by comparing spectral activity in LID animals on days when those animals were given L-DOPA alone (to induce LID) or L-DOPA+ketamine (**Fig. 6**). In the L-DOPA alone conditions, animals received 4 successive saline injections (2-h interval between injections), and then received a 5^th^ injection of saline paired with L-DOPA. Neural activity was assessed during the L-DOPA on-state. In the L-DOPA+ketamine condition, animals received 4 successive injections of ketamine alone (2-h interval between injections) and were given L-DOPA+ketamine on the 5^th^ and final injection. We predicted that broadband gamma would be reduced in the L-DOPA+ketamine condition in M1 and the striatum. The results, however, were mixed as paired *t*-tests indicated that L-DOPA+ketamine resulted in an increase in M1 broadband-gamma (*t*(6)=2.24, *p*=0.03, *d*=0.41), a reduction of DMS gamma (*t*(6)=3.54, *p*=0.03, *d*=0.56) and NAc (*t*(6)=3.65, *p*=0.01, *d*=0.59), and no effect in the DLS (*t*(6)=0.98, *p*=0.13, *d*=0.11). Consequently, the effects of ketamine on gamma in LID animals appears to be region specific, with gamma suppression only occurring in the medial and ventral striatum. An analysis of mean spectral power following each of the 5 injections can be found in **Supplementary Fig. 4**.

**Figure 6.**
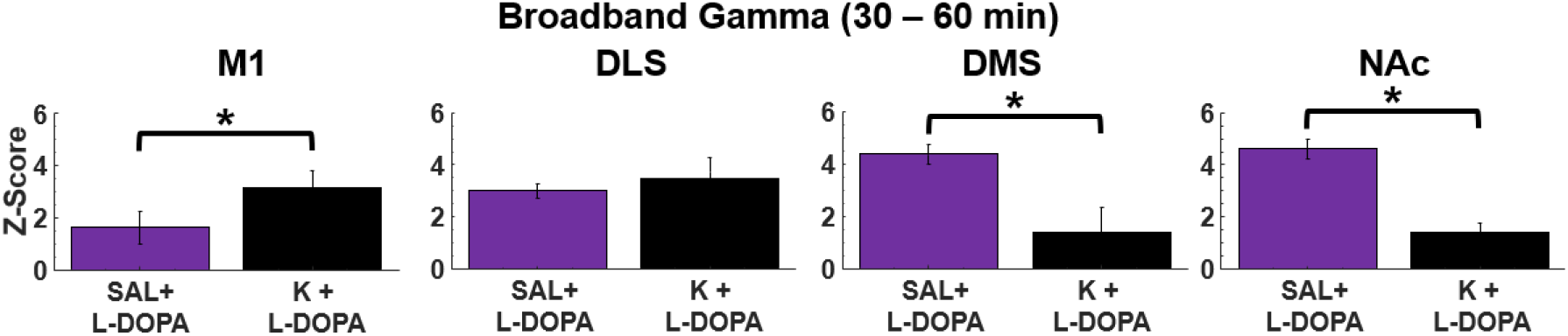
Ketamine increases L-DOPA-induced broadband gamma in M1 but reduced broadband gamma in the DMS and NAc. Injection of L-DOPA (7 mg/kg) after 21d priming with the same dose triggered broadband gamma oscillations (35-80 Hz, data from Figure 2E, purple bars, mean±SEM). Data was obtained 30-60 days post-L-DOPA priming. In the same animals (*N*=7), five subsequent injections of ketamine were administered with the 5^th^ injection paired with L-DOPA (7 mg/kg, data from Figure 4A/B, black bars). Paired t-test showed ketamine increased L-DOPA-induced broadband gamma in M1 (p=0.03) but reduced broadband gamma in the DMS (*p*=0.01) and NAc (*p*=0.01).

After investigating the hypothesis that ketamine reduces broadband gamma, we performed exploratory analyses to identify potential relationships between ketamine+L-DOPA on other frequency bands. For example, and as suggested in **Fig. 5**, a notably large increase in ∼140-Hz HFOs in DLS, DMS, and NAc was observed when ketamine was combined with L-DOPA during Injection 5 (ANOVA, all *p*<0.05, see **Supplementary Fig. 3** and **Supplementary Table 2**). It is well known that ketamine without L-DOPA induces robust HFOs (Nicolás et al., 2011; Caixeta et al., 2013, Ye et al., 2018). Consequently, L-DOPA may enhance coordination between neurons in circuits involved in generating ketamine-induced HFOs.

### Previous exposure to ketamine is associated with increased M1 broadband activity during LID

While it is clear that ketamine induces profound changes in the local-field signal within minutes of injection, a single 10-hour exposure to ketamine can reduce LID in rodent models for weeks (Bartlett et al., 2016). This suggests that ketamine induces lasting neuroplastic changes. Consequently, we investigated whether prior exposure (days) to ketamine alters spectral activity during the L-DOPA on-state. Spectral activity was analyzed during the 30-60-minute interval following the SAL+L-DOPA injection. Within-subject comparisons (*n*=7) were made between sessions in which rats had either no prior exposure to ketamine and received only saline+L-DOPA (“0 K” in **Fig. 7**) to sessions when animals had previously received at least one exposure to ketamine (“>1 K” in **Fig. 7**). Since the acute effects of ketamine were investigated in **Fig. 6**, sessions in which ketamine was also administered with L-DOPA were not included in this analysis.

**Figure 7.**
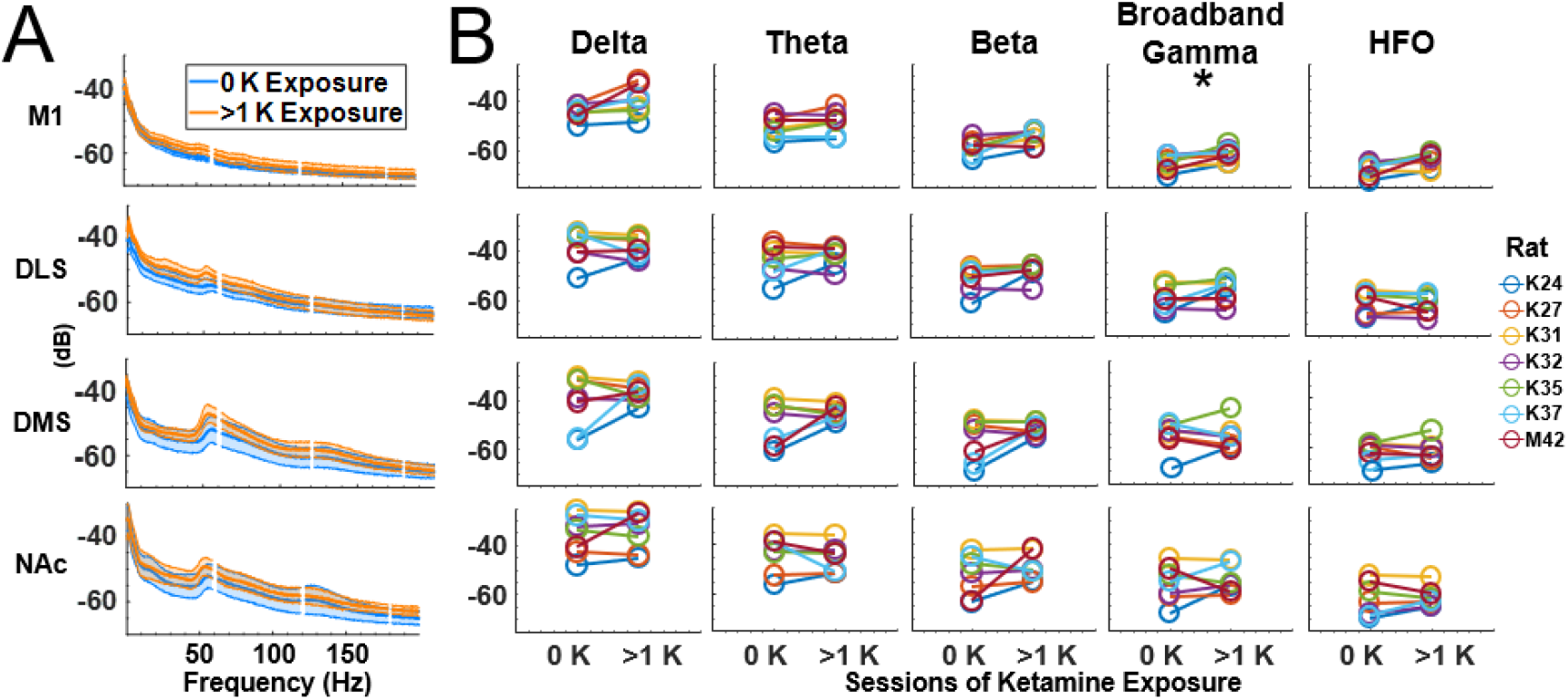
Prior ketamine exposure is associated with increased M1 broadband activity during LID. **(A)** Average power spectra (dB; mean±SEM) following L-DOPA exposure (30-60 min post injection) before any prior exposure to ketamine (blue) compared to data from sessions following at least one ketamine exposure session (orange). **(B)** Within-subject comparisons (paired *t*-tests, Holm-corrected) revealed significant long-term increases in broadband gamma in M1 (*p*=0.03). No other significant differences were observed.

Average spectral power (dB) for each region following L-DOPA administration is presented in **Fig. 7A**. Power-spectral responses following L-DOPA injections prior to ketamine exposure (blue) were compared to spectral responses that occurred during the weeks that followed ketamine exposures (orange). Contrary to the hypothesis that sustained ketamine exposure reduces broadband gamma power, Holm-corrected paired *t*-tests identified an increase in spectral power in broadband gamma in M1 (*p*=0.03, *d*=1.59). No significant effects were observed in the striatum. Given that it was not feasible to have an additional group of animals that was only injected with saline for the 4 weeks of neural recording for comparison, it is conceivable that this effect relates to some factor associated with the passage of time and not to ketamine. Even so, the fact that broadband gamma was not suppressed in this longitudinal experiment indicates that the therapeutic effects of ketamine are not a result of lasting gamma suppression during the L-DOPA on-state.

The preceding analysis examined the effect of prior exposure to ketamine on raw spectral power during LID. Alternatively, it is conceivable that prior ketamine exposure affects the change in the LFP signal induced by L-DOPA injection. To address this, we computed the baseline-subtracted power-spectral densities and performed the same analysis as described for **Fig. 7** (**Supplementary Figure 5**). This analysis identified no effect in any region or frequency band, further indicating that ketamine does not produce a lasting spectral signature in the striatum after it is metabolized. It also suggests that the increased M1 raw (not baseline subtracted) broadband gamma observed in **Fig. 7B** indicates persistently elevated broadband gamma that does not change following L-DOPA injection.

### Ketamine suppresses L-DOPA-induced theta-to-high-gamma corticostriatal CFC

Theta-to-high-gamma PAC was investigated since reduced theta-to-high-gamma PAC is a feature of LID in animal models (Belić et al., 2016). Given ketamine’s capacity to reduce LID, we hypothesized that ketamine administration would reduce PAC in LID rats. Theta-to-high-gamma PAC in the LID on-state (LID L-DOPA, 7 mg/kg) was compared to the condition when LID animals received ketamine (20 mg/kg) + L-DOPA (7 mg/kg, LID+K+L-DOPA) after 21 days of priming at 7 mg/kg (same data as in **Fig. 5**,**6**). The time-course of theta-to-high-gamma PAC is presented in **Fig. 8B** (right), and group-level comparisons are presented in **Fig. 9** (right column). Inspection of the time-course of theta-to-high-gamma PAC in the LID+L-DOPA group indicated that PAC increased in M1 and the striatum relative to the pre-injection baseline (**Fig. 8B**, red lines; **Fig. 9**, red bars), and was not apparent in the LID+K+L-DOPA group (**Fig. 8**, green lines; **Fig. 9**, green bars). Statistical analysis of PAC in **Fig. 9** (right column) was performed for the 30-60 min post-injection period as this is when L-DOPA reaches peak effect. ANOVA for the 5 conditions was performed for each brain region. Main effects of condition were identified in M1, DMS, and NAc (*F*_*M1*_(4,28)=6.3, *p*=0.001, η^2^=0.51; *F*_*DMS*_(4,31)=7.12, *p*=0.0004, η^2^=0.51; *F*_*NAc*_(4,28)=14.8, *p*=0.0001, η^2^=0.68), but not DLS (*p*>0.05).

**Figure 8.**
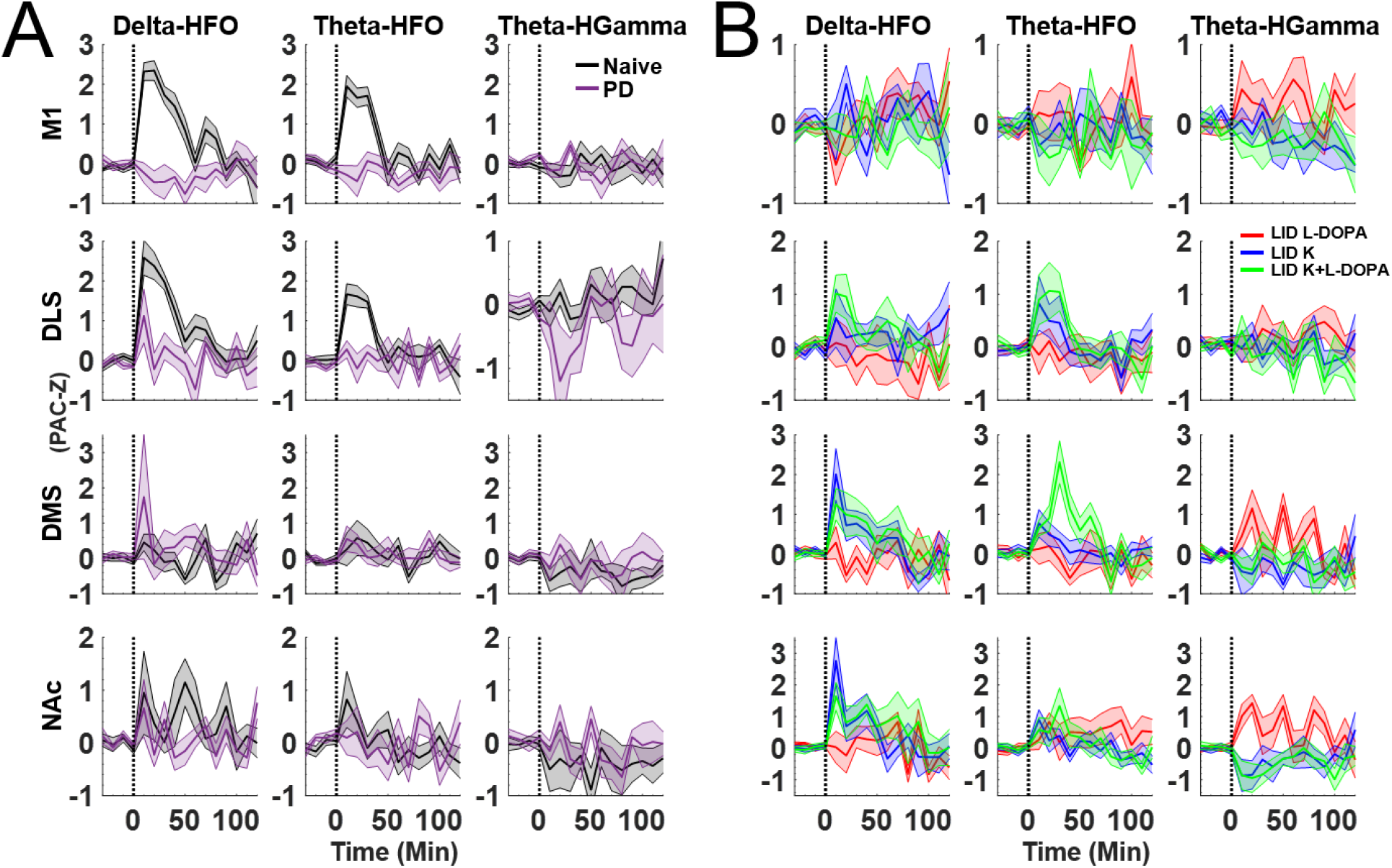
Ketamine induced cross-frequency coupling in naïve, 6-OHDA-lesioned, and LID animals. **(A)** Time course of drug-induced CFC by region. Data shown are averages (mean±SEM) of the 4^th^ injection of ketamine in naïve (black) and 6-OHDA-lesioned (purple) animals. **(B)** As in (A) but for LID animals receiving injections of L-DOPA (7 mg/kg) alone (red), ketamine alone (blue), or ketamine + L-DOPA (green). Data is from Figure 5 (21-d primed animals), recorded 30-60 days post-L-DOPA priming.

**Figure 9.**
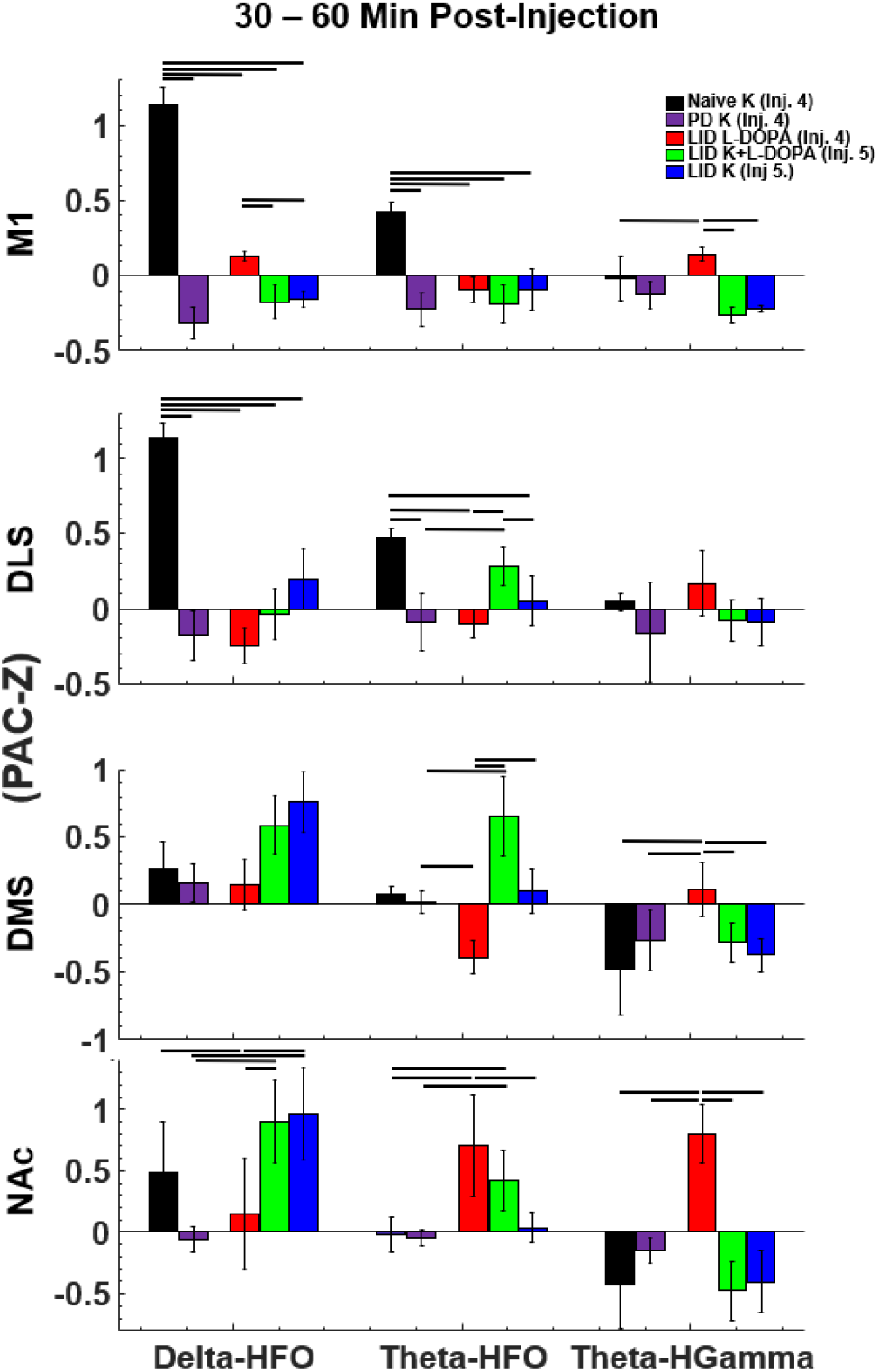
Ketamine suppresses cross-frequency coupling associated with LID. Data from Fig. 8 averaged (mean±SEM) between 30-60 min post-drug injection. ANOVA used for all statistics with Tukey-Kramer post-hoc correction. In the M1, the 4^th^ injection of ketamine in naïve animals triggered significant increases in delta-HFO and theta-HFO CFC (all *p*<0.05). In LID animals, co-injection of L-DOPA (7 mg/kg) + ketamine (green) significantly reduced delta-HFO coupling compared to L-DOPA alone (red) (*p*=0.04). Theta-high gamma CFC was significantly reduced in the motor cortex, DMS, and NAc following an injection of ketamine in naïve animals (black) and in LID animals (blue) even with L-DOPA on board (green) (all *p*<0.05). This suggests that ketamine selectively reduces CFC in the on-state of LID animals.

Theta-to-high-gamma PAC in LID animals following L-DOPA administration (red bars) was significantly reduced after co-administration with ketamine (green bars) in M1, DMS, and NAc (Tukey *post-hoc* corrected: *p*_M1_=0.02, *d*=1.56, *p*_*DMS*_=0.01, *d*=2.39, p_NAc_=0.0005, *d*=4.54), but not DLS (*p*>0.05). This effect was also observed when ketamine was delivered without L-DOPA in M1, DMS, and NAc (blue bars, LID off state; *p*_M1_=0.02, *d*=1.56, *p*_*DMS*_=0.01, *d*=2.39, p_NAc_=0.0005, *d*=4.54), but not DLS (*p*>0.05). We further explored ketamine’s effect on CFC in the 7-day primed animals (12 mg/kg L-DOPA, conditions as in **Fig. 2D**) and also observed a reduction in theta-to-high-gamma PAC, but only in the DMS (Bonferroni-Holm-corrected t-test, *p*=0.03; **Supplementary Fig. 6**). Taken together, these data suggest that ketamine impacts the oscillatory signature of LID by altering cross-frequency interactions and, specifically, by reducing theta-to-high-gamma PAC.

### Ketamine induces delta-HFO and theta-HFO cross-frequency coupling in naïve but not in PD and LID animals

Cross-frequency coupling is believed to support neural communication and neural plasticity by organizing the timing of action potentials (Canolty and Knight, 2010; Lisman and Jensen, 2013). Excessive CFC can also indicate circuit dysfunction. For example, increased theta- and delta-to-high-gamma phase-amplitude coupling in motor cortex is a signature of PD in primate models (Devergnas et al., 2019). Because acute exposure to ketamine is known to produce strong delta-HFO and theta-HFO coupling in the cortex and striatum (Cordon et al., 2015), we investigated whether acute exposure to ketamine alters CFC in naïve, PD, and LID hemi-lesioned rats.

In agreement with previous reports (Ye et al., 2018), ketamine injections produced robust delta- and theta-to-HFO PAC in M1 and striatum of naïve rats (**Fig. 8** and black bars in **Fig. 9**). Surprisingly, ketamine-induced delta- and theta-HFO CFC was absent in PD and LID rats (**Fig. 9:** black bars compared to PD and LID conditions). ANOVA identified a main effect of experimental condition (*e*.*g*., naïve, PD, LID) and CFC frequency band in M1 and DLS (M1: delta-HFO: *F*(4,28)=12.2, *p*=0.0001, η^2^=0.95; theta-HFO: *F*(4,28)=32.0, *p*=0.0002, η^2^=0.84; DLS: delta-HFO: *F*(4,31)=29.6, *p*=0.0001, η^2^=0.81; theta-HFO PAC: *F*(4,31)=13.82, *p*=0.0002, η^2^=0.67; NAc: delta-HFO: *F*(4,31)=15.36, *p*=0.001, η^2^=0.75). *Post-hoc* comparisons indicated that increased delta-HFO PAC in naïve (M1, DLS, and NAc, all *p*<0.001) but not PD/LID animals. The absence of coupling with HFOs in LID was surprising as HFO, theta, and delta power was high following each ketamine injection in the LID animals (**Fig. 5**). These results suggest that the low- and high-frequency oscillators activated by ketamine exposure become decoupled after prolonged dopamine depletion.

## Discussion

L-DOPA-induced dyskinesias are a debilitating consequence of dopamine-replacement therapy for PD. Although there is an abundance of research into the cellular and synaptic origins of LID, far less is known about how LID impacts systems-level circuits and neural synchrony. We investigated the oscillatory signatures of LID and explored how two commonly used LID priming procedures and L-DOPA dosages used to induce LID affect these signatures. Our first observation was that the dosage of L-DOPA used to induce LID impacts neural synchrony. We discovered that while 12 mg/kg L-DOPA delivered after the 7-day 12 mg/kg priming procedure induced focal corticostriatal 80-Hz gamma (Dupre et al., 2016; Halje et al., 2012), 7 mg/kg L-DOPA did not, yet both doses triggered robust LID. Furthermore, we found that non-oscillatory broadband gamma in the striatum and theta-to-high-gamma CFC in M1 and NAc was evident when the 21-day low-dose (7 mg/kg) procedure was used. We then explored how this activity was affected by ketamine, given evidence that ketamine reduces LID (Bartlett et al., 2016). We found that ketamine exposure during the LID on-state suppressed LID-associated striatal broadband gamma, enhanced broadband gamma in M1, and suppressed theta-to-high-gamma CFC in M1 and NAc. These data suggest that ketamine’s potential therapeutic effects are region specific. An unexpected finding was that while ketamine is known to induce robust theta- and delta-HFO CFC in naïve rats (Caixeta et al., 2013; Cordon et al., 2015; Ye et al., 2018), no such coupling was observed in PD or LID animals, despite these animals exhibiting robust HFOs. This suggests that prolonged dopamine depletion decouples neuronal networks involved in cross-band synchrony.

### L-DOPA dose determines the spectral signature of LID

While low-dose L-DOPA (6-7 mg/kg) is commonly used for LID priming and induction (Cenci et al., 1998; Dekundy et al., 2007; Bartlett et al., 2016), LID-associated oscillatory activity has only been investigated using high-doses of L-DOPA (12 mg/kg) (Dupre et al., 2016; Halje et al., 2012; Tamte et al., 2016). Using high-dose priming and induction, we replicated previous reports that LID is accompanied by focal 80-Hz oscillations in M1, DMS, and DLS (Dupre et al., 2016; Halje et al., 2012; Tamte et al., 2016). Surprisingly, these 80-Hz oscillations were absent when the 7 mg/kg L-DOPA dose was given to induced LID in these animals, despite this dosage producing clear behavioral evidence of LID (**Fig. 2**,**4**).

We then investigated how the low-dose, long-duration (7 mg/kg, 21-day) priming procedure affected synchrony. As will be discussed below, we found that these animals expressed theta-to-high-gamma CFC in M1 and NAc and non-oscillatory broadband gamma activity in the striatum during the LID on-state. Thus, L-DOPA can induce fundamentally distinct spectral states in animals expressing LID that depend on the dose and duration of exposure.

### Focal 80-Hz gamma following high-dose L-DOPA priming

Focal 80-Hz oscillations suggest tight temporal coordination between local networks of coupled inhibitory neurons (I-I) (Brunel and Wang, 2003; Buzsáki and Wang, 2012). In animal models, 80-Hz oscillations and L-DOPA-induced dyskinesias are reduced following the application dopamine receptor 1 (D1R) antagonists to the cortical surface (Halje et al., 2012). This suggests that activation of cortical D1 receptors can trigger and/or sustain these oscillations. D1 receptors (Towers and Hestrin, 2008) and *N*-methyl-D-aspartate receptors (NMDAR) (Lim et al., 2014) are expressed on M1 GABAergic interneurons and striatal medium spiny neurons (MSNs) which may partly explain why D1R and NMDAR antagonists interfere with these oscillations (Kirli et al., 2014) and why NMDAR antagonists reduce LID in human patients (Goetz et al., 2005; Oertel et al., 2017). As with Tamte et al. (2016), we observed 80-Hz gamma outside of M1 and specifically in the DMS. This suggests that 80-Hz gamma is either transmitted to DMS from M1 (Richter et al., 2013) or generated locally within the DMS. The latter possibility is conceivable as >95% of striatal neurons are inhibitory and tightly coupled and so, like M1, may be capable of sustaining high-frequency gamma oscillations. Finally, it is plausible that focal gamma is mediated by dopamine D4 receptors in the basal ganglia (Bello et al., 2019) as D4R-mediated gamma oscillations have been observed in the hippocampus (Andersson et al., 2012).

### Broadband gamma in LID emerges after weeks of low-dose priming

While clear 80-Hz oscillations emerged after 10 days of high-dose L-DOPA administration, no such activity was present when these animals were given the low L-DOPA induction dose (7 mg/kg). Instead, extended weeks-long low-dose priming in a separate group of rats resulted in broadband desynchronized activity. Unlike 80-Hz gamma, this broadband gamma was present in the striatum but not M1 (**Fig. 2**,**3**). To our knowledge, this is the first report of a new LID spectral signature using the low-dose 21-d priming procedure. The development of broadband gamma with repeated L-DOPA exposure suggests neuroplastic changes resulting from cycling dopamine levels (Calabresi et al., 2015) that may drive the reorganization of striatal circuits through the activities of fast-spiking interneurons (Berke, 2011). Similar high-frequency broadband activity has been interpreted as increased action-potential firing or an altered balance between excitation and inhibition (Rubenstein and Merzenich, 2003; Voytek et al., 2015). These ideas are consistent with the observation that L-DOPA treatment increases direct pathway activity (Albin et al., 1989; DeLong, 1990), and because optogenetic and chemogenetic manipulations that increase striatal direct-pathway activity produce LID in animal models (Alcacer et al., 2017; Perez et al., 2017; Rothwell et al., 2015). How this excitability emerges is an open question. It is conceivable that persistent L-DOPA treatment and chronically elevated dopamine levels increase dopamine-mediated plasticity in the absence of sensorimotor input (Fieblinger et al., 2014; Picconi et al., 2003; Shen et al., 2008). Striatal plasticity in the absence of input from the cortex could disrupt the mapping of striatal responses to afferents and result in less organized broadband activity. Furthermore, it is important to highlight that the 21-d (7 mg/kg) primed animals did have overall lower LID compared to the 7-d (12 mg/kg) (**Fig. 3B**). It is important to note that our top-down tracking of 2-D location may have missed behaviors such as rearing and grooming that could be enhanced or suppressed in LID animals and conceivably induce broadband gamma activity. This seems unlikely however as we did not observe any difference in overall movement between the 7-d and 21-d groups (*t*-test; *p*=0.09).

### LID-associated broadband gamma was reduced when L-DOPA was co-administration with ketamine

Given evidence that ketamine can reduce dyskinesias by ∼50% regardless of L-DOPA dose, 6-7 mg/kg and 12 mg/kg (Bartlett et al., 2020, 2016), we predicted that spectral signatures of LID would decrease after ketamine administration. Specifically, we hypothesized that ketamine administration would reduce the striatal broadband gamma activity we observed during LID in animals exposed to the 21-day low-dose priming procedure. In agreement, broadband gamma was reduced in DMS and NAc in the L-DOPA+ketamine condition relative to the L-DOPA alone condition (**Fig. 6**). Interestingly, suppression was not observed in DLS (no change) and M1 (increased broadband). Broadband gamma is linked to several physiological processes (Buzsáki and Wang, 2012) including enhanced neural excitability (Fries et al., 2007). L-DOPA treatment is believed to increase direct pathway activity (Albin et al., 1989; DeLong, 1990), and increased direct-pathway neural activity may be a component of the observed increase in broadband power in LID animal models (Alcacer et al., 2017; Perez et al., 2017; Rothwell et al., 2015). Consequently, it is conceivable that ketamine reduces pathological activation of the direct pathway to exert anti-dyskinetic effects. It is less clear why this effect is most prominent in NAc and DMS, but not DLS.

The ketamine-alone and ketamine+L-DOPA conditions all resulted in increased M1 broadband gamma (**Fig. 6**). Similar responses to ketamine and other NMDAR antagonists have been reported (Nicolás et al., 2011; Hunt and Kasicki, 2013, Ye et al., 2018) (**Fig. 5**). Gamma-generation in the cortex by ketamine could result from its antagonism of NMDARs on parvalbumin-expressing (PV) GABAergic interneurons as these NMDARs have a high affinity for ketamine (Hunt and Kasicki, 2013), and since it was shown that GABAergic interneurons, including PV-interneurons, but not glutamate principle neurons in the medial prefrontal cortex (mPFC), are the cellular trigger for ketamine’s rapid antidepressant actions (Gerhard et al., 2020). Antagonism of NMDARs has been proposed to reduce PV-neuron activity and consequently, disinhibit M1 principal cells (Buzsáki and Wang, 2012; Homayoun and Moghaddam, 2007; Hudson et al., 2020; Korotkova et al., 2010; Pinault, 2008). This could engage local networks of non-parvalbumin inhibitory cells contributing to gamma generation (Whittington et al., 1995), such as somatostatin interneurons that, in the mPFC, contribute to the antidepressive action of ketamine (Gerhardt et al., 2020). Moreover, patient and preclinical models of LID have shown increased glutamatergic activity (Oh et al., 1998; Chase and Oh, 2000; Calon et al., 2002). The combined effect of ketamine+L-DOPA on glutamatergic neurons may partially account for enhanced oscillatory activity. Dopamine may also be involved as ketamine-induced locomotion and gamma in M1 were eliminated when ketamine was delivered after administration of a D1R antagonist (Ye et al., 2018).

### Ketamine exposure is associated with a days-long increase in LID on-state broadband activity in the motor cortex

Ketamine exposure can induce neuroplastic changes that endure for weeks following a single exposure in rats (Li et al., 2010). Furthermore, a single 10-hour ketamine exposure can produce a weeks-long reduction in an animal model of LID (Bartlett et al., 2020, 2016), suggesting that ketamine-induced neuroplastic changes contribute to reduced LID. To investigate this, we examined LID on-state spectral activity for the days prior to and following the first exposure to ketamine. We found that broadband activity in M1 was increased in the days that followed the first 10-hour exposure to ketamine (**Fig. 7**). A limitation of our study is that L-DOPA-induced AIMs were not scored during the recording period of the 21-d priming groups. Consequently, we cannot directly correlate within-session broadband activity with LID severity. However, we did score LID during neurophysiological recordings when the 12 mg/kg L-DOPA induction dose was given on day 38 (**Fig. 2E**), and we observed that ketamine delivered 90 minutes later significantly reduced AIMs. While it is unclear why prior exposure to ketamine selectively increased broadband gamma in M1 but not the striatum, it is conceivable that differentially increased synaptogenesis and brain-derived neurotrophic factor (BDNF) production (Phoumthipphavong et al., 2016; Yang et al., 2013) in the cortex are responsible. Furthermore, the highly-selective NMDA receptor antagonist MK-801, a ligand with similar mechanisms to ketamine, has been found to increase gamma in the cortex by targeting pyramidal neurons and PV-interneurons (Hudson et al., 2020). Ketamine’s therapeutic effects also extend to alleviating amblyopia via inhibitory cortical PV-neurons (Grieco et al., 2020). Prior work has shown that the long-term anti-dyskinetic effect of ketamine correlates with changes in dendritic spine density, specifically a reduction of multi-synaptic mushroom spines, in the striatum, yet changes in dendritic spines in the M1 had not been investigated in that study. And this effect of cortically-derived BDNF release can be blocked by systemically inhibiting BDNF-signaling (Bartlett et al., 2020).

### L-DOPA-induced theta-to-high-gamma CFC in PD and LID rats is suppressed by ketamine

In healthy animals, CFC may facilitate information transfer between brain regions (Canolty and Knight, 2010) or organize the timing of neural ensemble activity to support learning and memory (Lisman and Jensen, 2013). The potential roles of CFC in LID are largely unexplored, and only one study has investigated CFC in LID using the high-dose condition (Belić et al., 2016). This study identified reduced theta-to-high-gamma coupling in LID animals despite these animals expressing strong 80-Hz oscillations. Like Belić et al., we hypothesized that theta-to-high-gamma activity would be suppressed in LID rats, and that the novel broadband gamma activity we observed in LID under 7 mg/kg priming would be decoupled from low-frequency oscillations. Instead, we observed that theta-to-high-gamma CFC increased in M1, DLS, and NAc of LID animals (**Fig. 9**), suggesting that the priming procedure and dose can trigger distinct oscillatory neural states that are not a simple linear function of L-DOPA dosage. We also observed a significant increase in theta-to-high-gamma CFC in the DMS of the 7d prime group but only after 1 month without L-DOPA exposure (**Supplementary Fig. 6A**). A single ketamine injection delivered 90 minutes after L-DOPA significantly reduced this effect (**Supplementary Fig. 6B**), and further supports the argument that theta-to-high-gamma PAC in the DMS is associated with LID and suppressed by ketamine. Regarding broadband gamma, we did not observe theta-to-broadband gamma CFC in LID animals primed for 21+ days with 7 mg/kg L-DOPA, suggesting that theta-to-broadband gamma CFC is not a signature of LID.

### Ketamine enhances delta-HFO and theta-HFO cross-frequency coupling in naïve but not PD and LID animals

Multiple groups have reported that sub-anesthetic exposure to ketamine and other NMDAR antagonists induces strong HFOs (120–160 Hz) in the striatum (Olszewski *et al*., 2013; Cordon *et al*., 2015; Hunt *et al*., 2015, Ye *et al*., 2018), cortex (Cordon et al., 2015; Nicolás et al., 2011), and hippocampus (Caixeta et al., 2013). Some have also reported that these HFOs are coupled to the phase of delta and theta oscillations (Caixeta et al., 2013; Cordon et al., 2015, Ye et al., 2018). In support of this, we observed robust ketamine-induced HFOs in M1 and the striatum of naïve, PD, and LID rats (**Fig. 5**). Surprisingly, while delta- and theta-HFO CFC was clear in naïve rats, such coupling was not present in the PD or LID animals (**Fig. 9**). This suggests that prolonged dopamine depletion reconfigures circuits involved in synchronizing low-frequency oscillations with HFOs. It is conceivable that decoupling between delta/theta oscillations and HFOs is a physiological indicator of PD progression similar to beta-band synchrony (Brown, 2007). Future studies are required to determine if ketamine-induced delta- and theta-HFO CFC changes with disease progression.

## Conclusions

Our findings provide new insights and a more nuanced view of how PD and LID impact neural coordination in corticostriatal circuits. While we replicated the observation that focal 80-Hz oscillations are a robust signature of LID, we also found that this signature depends on the dose of L-DOPA administered, where low-dose induction results in clear LID but without 80-Hz gamma. We also found that striatal broadband gamma and corticostriatal theta-to-high-gamma CFC were enhanced in LID rats under the common 7 mg/kg, 21-day procedure. These differences suggest that the neural signatures of L-DOPA-induced dyskinesia do not lie on a continuum that depends on dose, but that the dose can produce distinct threshold-dependent neuronal states in corticostriatal circuits. Ketamine was also found to reduce the enhanced CFC observed in LID under the low-dose priming protocol, suggesting that a component of ketamine’s anti-dyskinetic effect is through suppression of theta-to-high-gamma coupling. These results add to growing evidence for the mechanistic basis of ketamine’s therapeutic effects and complement ongoing clinical testing of sub-anesthetic ketamine for the treatment of LID by our group.

## Acknowledgements

We would like to give special thanks to Matthew Schmit, Paulluvi Bahl, Maddy Otto, and the collective undergraduates of the Cowen and Falk-Sherman laboratories for their contributions. MJB would like to thank the Society for Neuroscience’s Neuroscience Scholars Program for his Fellowship.

